# Compressive peripheral processing and across-channel suppression models of loudness bandwidth effects

**DOI:** 10.64898/2026.11.27.769709

**Authors:** Yi Shen, Stephen T. Neely

## Abstract

Band-limited sounds can be quieter than equal-level tones, contrary to classical loudness summation. Categorical loudness scaling (CLS) by normal-hearing listeners for flat noise (*N* = 30) and amplitude-modulated tones (*N* = 20) at 250 Hz to 4 kHz was combined with published loudness matches for wider complexes (*N* = 7) to compare nine central read-outs. Peripheral compression alone failed on wideband noise. Five across-channel suppression models reached the 1.68-dB listener reference and were indistinguishable on CLS. Global neural-ensemble averaging was the most parsimonious, whereas divisive normalization gave the lowest error on the wider complexes without refitting (1.90 dB).

## 1. Introduction

Classical loudness models predict that spreading energy beyond one auditory-filter bandwidth increases loudness, because total loudness sums compressive specific loudness across characteristic frequency (Glasberg and Moore, 2002; Moore *et al*., 1997; Zwicker and Scharf, 1965). Behavioral studies, however, show that band-limited sounds are often quieter than equal-level pure tones. Adaptive loudness matching revealed 2 to 5 dB of mid-bandwidth loudness depression for subcritical multitone complexes and noise bands (Hots *et al*., 2014; Leibold *et al*., 2007). Single-channel nonlinear cochlear modeling showed that peripheral compression and two-tone suppression produce only 0.5 to 2 dB of reduction for subcritical noise, far below behavioral depression, pointing toward retrocochlear mechanisms (Hots *et al*., 2016). Multi-frequency categorical loudness scaling (CLS) likewise showed negative bandwidth effects across 250 Hz to 4 kHz for 1/4- and 1-octave flat noise (Neely *et al*., 2026). The current study evaluates which phenomenological peripheral and central operations parsimoniously account for these departures.

Two mechanisms could underlie negative bandwidth effects. First, within-channel envelope fluctuations from auditory filtering interact with a memoryless compressive nonlinearity (via Jensen’s inequality, attenuating peaks relative to an equal root-mean-square [RMS] tone) or central modulation processing (Dau *et al*., 1997; Hots *et al*., 2016, 2014; Jepsen *et al*., 2008). Second, across-channel suppression, hypothesized to reflect brainstem or midbrain inhibitory pooling, reduces pooled activity as excitation spreads across channels (Carandini and Heeger, 2012; Oertel *et al*., 2011; Palmer *et al*., 1996; Pressnitzer *et al*., 2001; Rabinowitz *et al*., 2011; Winter and Palmer, 1995). Flat noise confounds spread with Rayleigh fluctuations, whereas sinusoidally amplitude-modulated (SAM) tones separate them: 20-Hz SAM (40-Hz bandwidth) isolates within-channel modulation, 100-Hz SAM (200-Hz bandwidth) transitions from resolved sidebands below 1.3 kHz to unresolved modulation above 1.3 kHz, and 1-octave SAM presents three resolved components. Because off-frequency summation cancels within-channel depression under classical summation, across-channel suppression is needed to unmask noise depression while keeping resolved 1-octave SAM near zero or weakly positive.

The current study had two aims: first, combining normal-hearing (NH) CLS data for flat noise and SAM tones (250 Hz to 4 kHz) with broadband matching data (Leibold *et al*., 2007) to separate modulation from spread; second, pairing a shared peripheral stage with nine central read-outs across four classes (classical summation, modulation, suppression, and hybrids) to evaluate accuracy, CLS parsimony, periphery sensitivity, and external broadband generalization.

## 2. Methods

### 2.1 Behavioral datasets and primary frequency range

Three NH datasets were compiled (`SuppPub2.xlsx`), including two CLS datasets using identical hardware (44.1 kHz, RME Babyface Pro, Etymotic ER-3A earphone) and MATLAB quick-Categorical Loudness Scaling (qCLS) software (Shen *et al*., 2024); 11-category ratings (0 to 50 categorical units [CU], ISO 16832:2006) were fitted with the five-component model (Shen *et al*., 2024) to derive equal-loudness contours. First, the Boys Town National Research Hospital (BTNRH) dataset (Neely, 2026; Neely *et al*., 2026) comprised 30 NH adults (19 to 69 years, mean 37.7, SD 15.3; pure-tone average [PTA; 0.5 to 2 kHz] ≤ 15 dB HL; better ear) completing two randomized 100-trial runs per condition for 1-s, 200-ms-ramped pure tones and 1/4- and 1-octave flat-spectrum noise (one frozen token per condition). Second, the University of Washington (UW) dataset enrolled 20 NH adults (18 to 28 years, mean 20.9, SD 2.3; 12 females, 8 males; PTA4 [0.5 to 4 kHz] ≤ 20 dB HL, interaural difference *<* 10 dB; test-ear mean 6.4, SD 4.4 dB HL) completing one 100-trial run per condition without practice (five randomized conditions) for 1-s, 200-ms-ramped tones and four 100%-modulated, RMS-normalized SAM tones (sine modulator on cosine carrier): 1/4-octave (*f_m_* = *f_c_*(2^1^*^/^*^4^ − 1)*/*(2^1^*^/^*^4^ + 1)), 1-octave (*f_m_* = *f_c_/*3), 20-Hz, and 100-Hz SAM.

All eight primary center frequencies (250, 500, 750, 1000, 1500, 2000, 3000, and 4000 Hz) were processed identically; data at 6 kHz were excluded because band stimuli were scaled by a single calibration value at the center frequency, which misstates the delivered level where the earphone response falls steeply (Sec. S3). Bandwidth or amplitude modulation (AM) influence was defined as equally loud pure-tone level minus stimulus level (dB) from 30 to 100 dB SPL in 10-dB steps (negative indicates depression), invariant to monotonic loudness-scale transformations. Excluding four 100-dB BTNRH endpoints outside the invertible tone range left 380 valid observations across 48 curves (62 each for BTNRH noise; 64 each for UW SAM). Per-listener UW 60-phon differences matched group-mean 60-dB SPL CU inversion closely across all 32 frequency-by-condition points (*r* = 0.968). A listenersampling SE reference (1.68 dB; 1.52 to 1.81 dB) pooled per-point bootstrap SEs across five conditions (four UW SAM conditions and BTNRH 1-octave noise; Neely *et al*., 2026 also reported a 1.80-dB RMS SE for 1/4-octave noise). Third, Leibold *et al*. (2007) group-mean adaptive matches (*N* = 7 NH adults; 300-ms, 10-ms-ramped, 1-kHz, 60-dB SPL five-tone complexes at 46 to 2119 Hz; SEs ≈ 1.1 to 1.8 dB) served as a single held-out external benchmark (unweighted RMSE across six bandwidths), excluded strictly from all fitting.

### 2.2 13-parameter peripheral stage with a memoryless compression nonlinearity

Stimuli (32 kHz, flat acoustics, 500 ms, 100-ms ramps; 10 frozen noise tokens) passed through 60 fourth-order complex gammatone filters spaced uniformly in equivalent rectangular bandwidth (ERB) rate *z* from 100 Hz to 10 kHz (Δ*z* = 0.547 ERB; Glasberg and Moore, 1990), giving envelopes *e_k_*(*t*). Gain-scaled level *L*_env_*_,k_*(*t*) = 20 log_10_[max(*G*(*f_k_*)*e_k_*(*t*)10*^L/^*^20^, 10*^−^*^12^)] passed through a memoryless sample-by-sample compression and rate nonlinearity (no temporal state beyond the filterbank) yielding excitation *E_k_*(*t*) and rate *s_k_*(*t*) (Neely and Jesteadt, 2005),

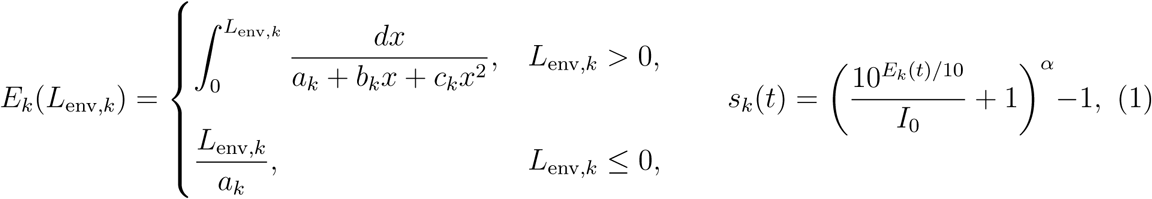

where *x* is a dummy variable, *a_k_*, *b_k_*, and *c_k_* are linear-in-*u* compression coefficients (*u* = log_10_(*f/*1000 Hz)), *I*_0_ is the intensity knee, and *α* is the rate exponent. Outer- and middle-ear gain log_10_ *G*(*u*) was a cubic polynomial clamped to *u* ∈ [−0.602, +0.602] (250 to 4000 Hz), giving 13 peripheral parameters including scale constant *C* (Table S1a). Time-averaged steady-state channel rate *s̄_k_* = ⟨*s_k_*(*t*)⟩*_t_* (0.12 to 0.38 s) drove static read-outs (*s_k_*(*t*) drove the modulation filter bank); 1-s, 30-token checks confirmed convergence (RMS difference = 0.14 dB; Sec. S2). Equal-loudness levels were inverted via log-loudness interpolation on a 1-dB tone grid (−10 to 120 dB SPL; Sec. S1). A five-way ablation replaced *e_k_*(*t*) with RMS envelopes before Eq. (1) (preserving tone growth), removed quadratic compression (*E_k_* = *L*_env_*_,k_*), linearized the rate stage, or linearized both (Table S4).

### 2.3 Candidate central read-out models and evaluation

Nine central read-outs across four classes (Table S1b) were evaluated on all 380 observations (250 Hz to 4 kHz), first on a shared 13-parameter pure-tone periphery (three alternation cycles around Eq. (3) constrained to BTNRH tone loudness growth; Sec. S1) and then with an independently matched pure-tone periphery per model. Every static read-out sums channels linearly, N = CΔz ∑_k_ r_k_, with no extra post-pooling exponent.

*Class 1 (Classical summation baseline):* Model M0 sets *r_k_* = *s̄_k_* (*k* = 0), isolating peripheral compression (Eq. (1)) without central suppression.

*Class 2 (Within-channel modulation processing):* Model M3a applies a 12-band modulation filter bank (MFB; *f_m_* = 5 to 1000 Hz, *Q* = 1.5; Dau *et al*., 1997) to *s_k_*(*t*) to compute Parseval-normalized modulation power *P_m,k_*, low-pass depth 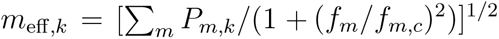, and attenuation 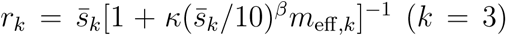. Separately, Dau *et al*. (1997) five-stage feedback adaptation loops (*τ* = 5 to 500 ms) replaced Eq. (1) on gammatone envelopes under classical linear summation at 1 kHz and 60 dB SPL.

*Class 3 (Across-channel spectral-spread suppression):* Five phenomenological models implement spectral-spread suppression via three mathematical structures. First, local subtractive wideband inhibition (local sub-WBI; Palmer *et al*., 1996; Winter and Palmer, 1995) subtracts a Gaussian-weighted local pool 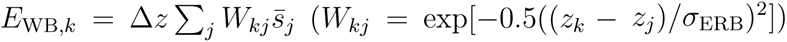 before rectification, *r_k_* = max(*s̄_k_* − *I*_WB_*_,k_,* 0), with linear *I*_WB_*_,k_* = *γE*_WB_*_,k_* (M2a, *k* = 2) or saturating 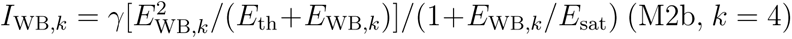. Second, neural-ensemble averaging (NEA; Neely *et al*., 2026) is a global, spectral-spread-dependent read-out (with no temporal state or adaptation filter) blending the broad sum *N*_sum_ and peak activity max*_k_ s̄_k_* (high-exponent limit of a peak-weighted average),

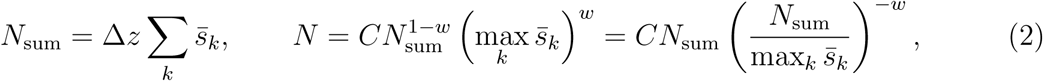

where *w* is the blend weight (*w* = 0 summation; *w* = 1 peak tracking). We evaluated global NEA with constant *w* = *w*_0_ (M1a, *k* = 1) or an internal-activity-gated Gaussian *w* = *w*_0_ exp(−[(log_10_ *N*_sum_ − *µ*_0_)*/σ_w_*]^2^) (M1b, *k* = 3); lacking a finite pooling width, Eq. (2) suppresses broad spectra regardless of component separation. Third, local divisive normalization (Carandini and Heeger, 2012) applies local shunting pooling within a Gaussian kernel of SD *σ*_ERB_ (M2c, *k* = 3),

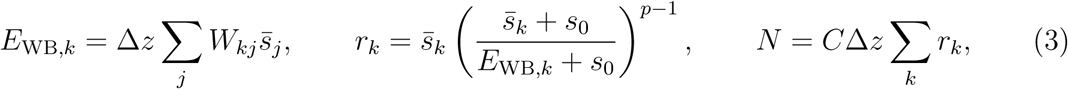

where *s*_0_ *>* 0 is the semi-saturation constant, *p* ≥ 1 is the normalization exponent, and *s̄_k_* and *E*_WB_*_,k_* are floored at 10*^−^*^12^. Intuitively, *s*_0_ preserves linear summation (*r_k_* → *s̄_k_*) near threshold, whereas above threshold *p* = 1 yields summation and *p* = 2 yields quadratic local normalization (*r_k_* ≈ *s̄*^2^*/E*_WB_*_,k_*) until bandwidth exceeds the local kernel span and broadband summation recovers.

*Class 4 (Hybrid modulation and spectral-spread suppression):* Six-parameter hybrids paired the MFB with Eq. (3) (M4b) or gated NEA (M5). CLS parsimony used BIC_eff_ = *n*_eff_ ln(RMSE^2^) + *k* ln(*n*_eff_), with primary *n*_eff_ = 48 curves (sensitivity checked across *n*_eff_ ∈ {12, 24, 30, 48, 380}; Table S3). Uncertainty (fixed shared periphery) used 48-curve cluster bootstrap, exact 20-listener UW resampling (*B* = 1, 000 on 256 points from `SuppPub2.xlsx`), CLS transfer, and eight-fold conditional central-read-out leave-one-frequency-out (LOFO) cross-validation (Sec. S2).

## 3. Results

### 3.1 Empirical dissociation across 250 Hz to 4 kHz

Figure 1 shows all 380 valid observations (250 Hz to 4 kHz). Pure-tone loudness growth agreed closely across laboratories (*r* = 0.996; MAE 1.16 CU at 30 to 80 dB SPL, 3.85 CU at 90 to 100 dB SPL). Subcritical 20-Hz SAM (third panel) showed consistent depression across frequency and level (mean −3.02 dB), isolating within-channel depression without spectral spread. The 100-Hz SAM condition (fourth panel) crossed at 1.3 kHz (dashed line), shifting from weak summation at ≤ 1 kHz (+0.36 dB) to deep depression at 1.5 to 4 kHz (−3.21 dB). Quarter-octave SAM (fifth panel) recovered from low-level depression toward 0 dB at higher levels, whereas resolved 1-octave SAM (rightmost panel) stayed modestly positive (+0.98 dB). Both BTNRH noise bands (first two panels) remained predominantly negative (−2.70 and −1.93 dB). Thus, within-channel modulation yields 3 to 6 dB of depression, whereas resolved spread gives weak summation for steady 1-octave SAM but net depression for fluctuating noise.

**Fig. 1.**
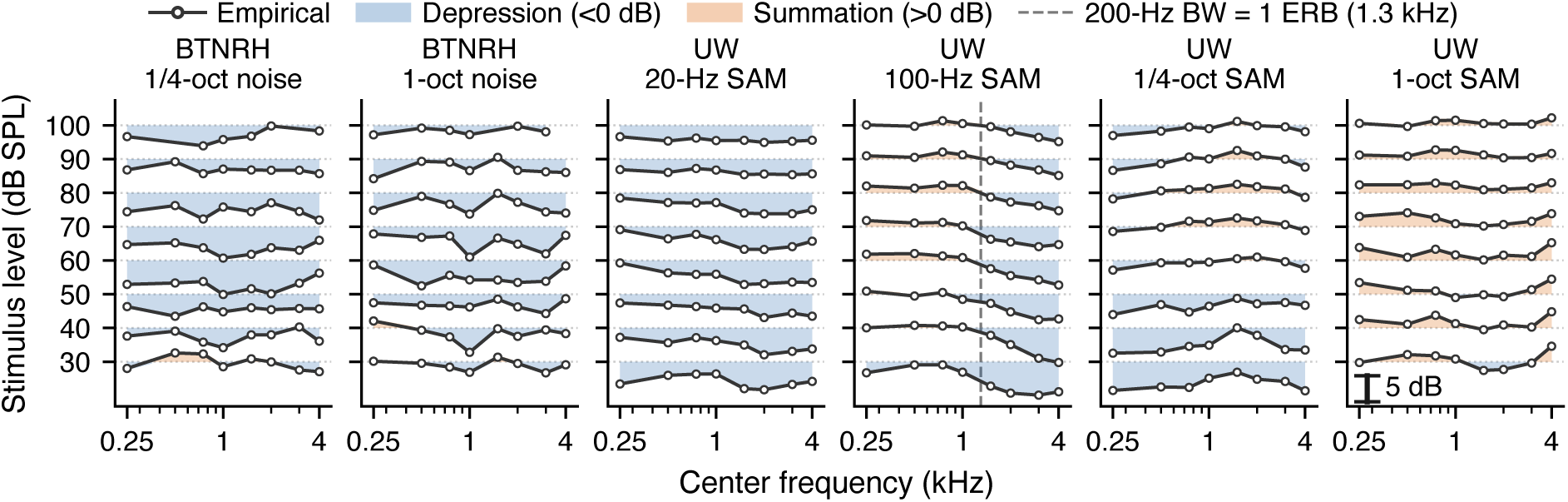
(Color online) Empirical bandwidth and AM effects across 250 Hz to 4 kHz (380 observations, 48 curves) plotted as offset level profiles (30 to 100 dB SPL, 10-dB steps). Left to right: BTNRH 1/4- and 1-octave noise (*N* = 30) and UW 20-Hz, 100-Hz, 1/4-octave, and 1-octave SAM tones (*N* = 20). Blue marks depression (*<* 0 dB), orange marks summation (*>* 0 dB), and the dashed line marks 1.3 kHz (200-Hz bandwidth ≈ 1 ERB); missing 100-dB points mark four excluded BTNRH endpoints, with a 5-dB scale bar at right.

### 3.2 Nine-model benchmark, peripheral ablation, and uncertainty

Figure 2 summarizes all nine models on 250 Hz to 4 kHz (Tables S1b to S3). On the shared 13-parameter periphery, Eq. (3) matched pure-tone growth to 1.85 dB RMSE; independently matched peripheries (diamonds, left panel of Fig. 2; 1.14 to 1.68 dB pure-tone RMSE, Sec. S1) left all competitors unchanged or worse.

**Fig. 2.**
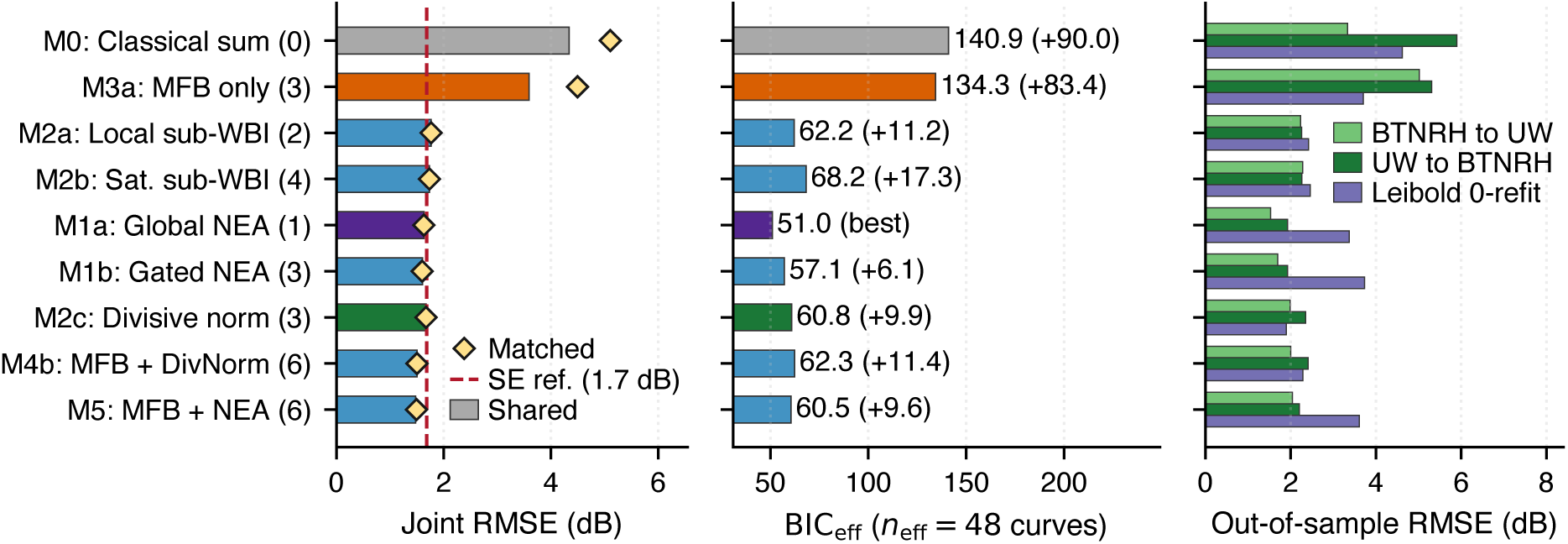
(Color online) Nine-model benchmark across 380 CLS observations (250 Hz to 4 kHz) and six external broadband conditions (Table S2). Left to right: joint RMSE, BIC_eff_ (*n*_eff_ = 48 curves), and out-of-sample RMSE. Left: shared-periphery bars (purple: M1a; green: M2c), matched peripheries (diamonds), and 1.7-dB SE reference (dashed). Right: CLS transfer (green) and zero-refit error on Leibold *et al*. (2007) (*N* = 7, purple).

Models without across-channel suppression (Classes 1 and 2) failed on wideband stimuli. Under classical summation (M0, joint RMSE = 4.34 dB), peripheral compression alone (Eq. (1)) predicted mean influences of −2.28 dB for 20-Hz SAM, −1.41 dB for unresolved 100-Hz SAM, and −2.83 dB for the 46-Hz Leibold *et al*. (2007) complex, but only −0.17 dB for 1/4-octave noise and strong overshoot on 1-octave noise (+5.44 dB). Five-way ablation confirmed instantaneous envelope compression is essential (Table S4): RMS pre-averaging eliminated subcritical depression (M0 20-Hz SAM = +0.20 dB; refitted M2c degraded from 1.67 to 2.43 dB), and removing either static nonlinearity reduced depression (M0 20-Hz SAM: −1.43 dB rate-only; −0.88 dB quadratic-only; +0.09 dB linear). Standalone MFB (M3a) reduced error modestly (3.59 dB), and Dau *et al*. (1997) feedback adaptation predicted the wrong sign on flat noise (+1.86 and +5.31 dB at 1 kHz, 60 dB SPL; Sec. S2).

Every Class 3 suppression model reached the 1.68-dB listener SE reference (1.61 to 1.76 dB; Fig. 2, Table S2). Crucially, CLS data alone did not separate the five Class 3 models: 48-curve cluster-bootstrap comparisons against M2c (1.67 dB; LOFO CV = 1.71 dB; M1a *w*_0_ = 0.636, LOFO CV = 1.64 dB) all spanned zero (M1a minus M2c = −0.04 dB, 95% CI [−0.14, +0.06]; Table S3). The exact 20-listener UW bootstrap (*B* = 1, 000; Table S5) confirmed M2c beat M0 in 100% of resamples (+1.61 dB) while overlapping all other Class 3 models (M1a minus M2c = +0.07 [−0.11, +0.30] dB). Under curve-level BIC_eff_ (*n*_eff_ = 48 and {12, 24, 30}) and CLS transfer, global NEA (M1a) ranked best (BIC_48_ = 51.0 vs. 60.8 for M2c and 62.2 for M2a), whereas point-level *n*_eff_ = 380 favored six-parameter hybrids (M5: 330.8, M4b: 345.4 vs. M1a: 378.7; Table S3). Adding the MFB (M4b, *k* = 6) lowered CLS RMSE by 0.17 dB while worsening BIC_48_ and broadband error (2.29 vs. 1.90 dB).

Model structures separated only on Leibold *et al*. (2007) (*N* = 7; Fig. 2, Table S5). Without refitting, M2c (*p* = 2.129, *σ*_ERB_ = 3.40 ERBs, *s*_0_ = 4.06; CIs in Table S1b) yielded the lowest zero-refit RMSE (1.90 dB; CLS parameter-bootstrap CI [1.67, 2.81]), beating global NEA in 99.1% to 99.6% of resamples (M1a: 3.37 dB; M1b: 3.73 dB) while overlapping local sub-WBI (M2a/M2b: 2.42 to 2.46 dB). One-parameter refitting lowered RMSE to 1.79 dB for M1a (*w*_0_ = 0.39) and 1.48 dB for M2c (*p* = 1.95).

Figure 3 overlays global NEA (Eq. (2)), divisive normalization (Eq. (3)), and M0 onto data. Whereas M0 diverges as bandwidth widens, both suppression models track mean levels within roughly one SE (M1a: 1.20 to 1.93 dB; M2c: 1.18 to 1.95 dB; UW 90 to 100 dB SPL residuals [1.03, 0.98 dB] were smaller than at 30 to 80 dB SPL [1.65, 1.82 dB]). Eq. (2) achieved lower RMSE on 100-Hz, 1/4-octave, and 1-octave UW SAM (100-Hz SAM *r* = +0.74 vs. +0.62; 1-octave SAM 1.20 vs. 1.53 dB, *r* = +0.09 vs. −0.18), whereas Eq. (3) achieved stronger BTNRH noise correlations (1/4-octave +0.58 vs. +0.12; 1-octave +0.38 vs. −0.03).

**Fig. 3.**
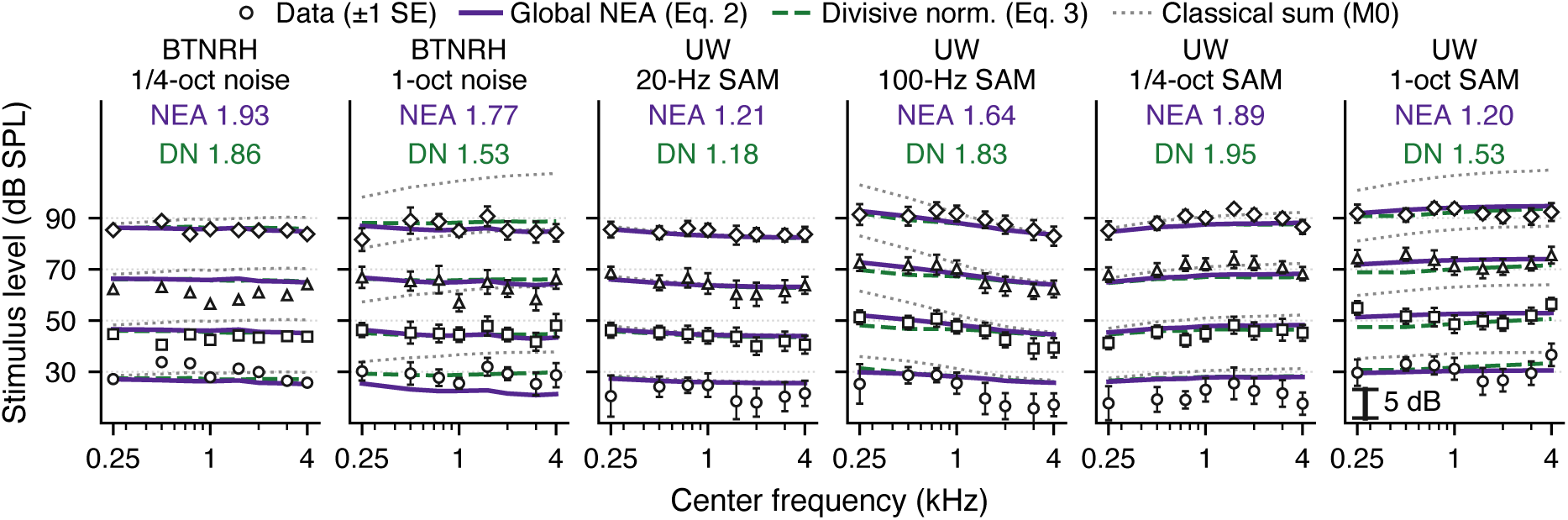
(Color online) Empirical data vs. global NEA (Eq. (2), solid purple), divisive normalization (Eq. (3), dashed green), and M0 (dotted gray) across 250 Hz to 4 kHz at 30, 50, 70, and 90 dB SPL (offset). Panels follow Fig. 1, with open symbols showing data (±1 SE where available); insets report all-level RMSE (dB) for NEA (purple) and DN (green), with a 5-dB scale bar at right.

### 3.3 Diagnostic predictions, level dependence, and external broadband check

Figure 4 compares global NEA (Eq. (2), solid) and divisive normalization (Eq. (3), dashed) across four diagnostic tests. At 60 dB SPL (far left), M0 overshoots 1-octave noise strongly, whereas both suppression models unmask compressive depression and reproduce the 2 to 3 dB noise-versus-SAM separation (Eq. (2) matching positive 1-octave SAM closer).

**Fig. 4.**
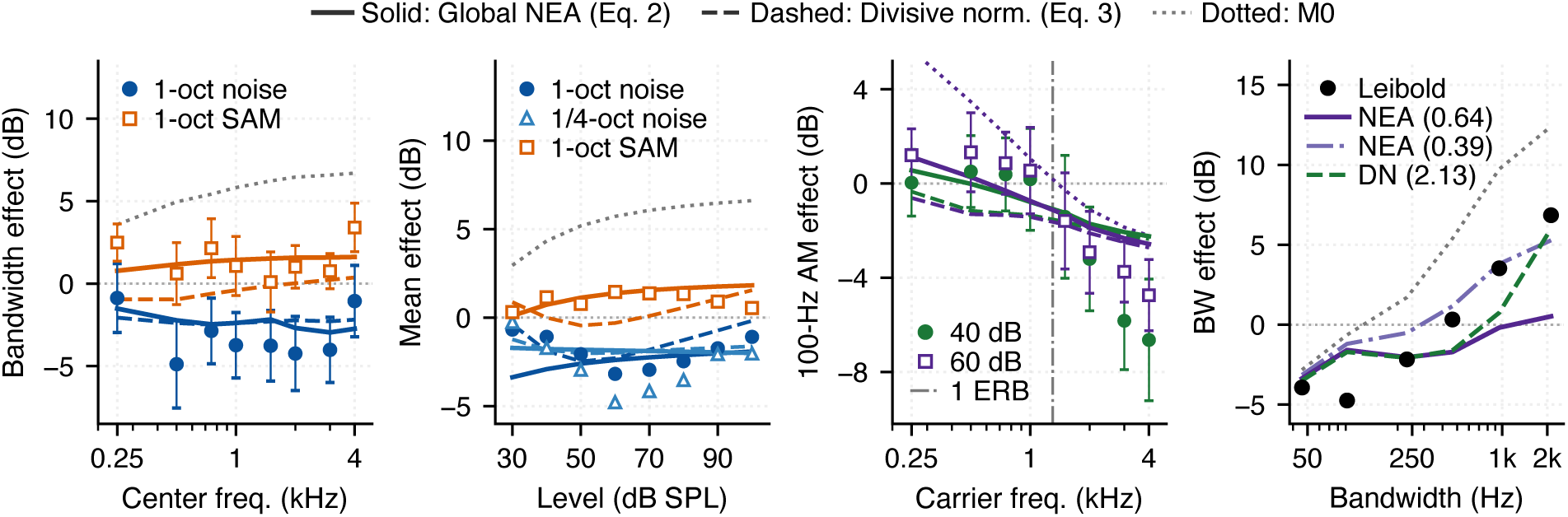
(Color online) Diagnostic comparison of global NEA (Eq. (2), solid), divisive normalization (Eq. (3), dashed), and M0 (dotted). Left to right: 1-octave noise vs. SAM (60 dB SPL), frequency-averaged level dependence (30 to 100 dB SPL), 100-Hz SAM 1-ERB crossover (40 and 60 dB SPL), and Leibold *et al*. (2007) (1 kHz, 60 dB SPL, *N* = 7). Symbols denote data (±1 SE where shown); far right shows zero-refit NEA (*w*_0_ = 0.64, solid purple), refitted NEA (*w*_0_ = 0.39, dash-dotted), and zero-refit DN (*p* = 2.13, dashed green).

Across 30 to 100 dB SPL (second panel, averaged over 250 Hz to 4 kHz), both noise bands show a U-shaped mid-level dip (1-octave: −0.7, −3.2, and −1.1 dB at 30, 60, and 100 dB; 1/4-octave: −0.3, −4.8, and −2.0 dB), whereas 1-octave SAM stays positive (+0.3 to +1.5 dB). Global NEA (solid) tracks 1-octave SAM closely (+0.1 to +1.8 dB) and recovers monotonically on 1-octave noise (−3.4 to −1.9 dB) via pure-tone broadening, but lacks threshold semi-saturation and misses the 30-dB return toward 0 dB. Divisive normalization (dashed) reproduces the full U-shape (−0.4, −2.5, −2.3, and −0.2 dB at 30, 50, 60, and 100 dB) via threshold linearity (10*^Ek/^*^10^ *< I*_0_, *s*_0_), shunting pooling, and high-level pure-tone broadening as filter skirts rise above *I*_0_ (unchanged within 0.00 dB at *c_k_* = 0; Sec. S2), though its 1/4-octave dip stays shallow (−1.8 vs. −4.8 dB at 60 dB). In the third panel, both models track the 100-Hz SAM 1-ERB crossover (1.3 kHz), with Eq. (2) capturing low-frequency positive summation better. At far right, zero-refit Eq. (2) (*w*_0_ = 0.64) captures subcritical depression but plateaus above 1 octave (3.37 dB; 1.79 dB refitted to *w*_0_ = 0.39), whereas zero-refit Eq. (3) (*p* = 2.13) restores summation above 1.5 octaves (1.90 dB; 1.48 dB refitted to *p* = 1.95; preserved under 300-ms randomized-phase checks, Sec. S2). Across 50 to 90 dB SPL, a +0.1 shift alters 1-octave matches by −1.37 to −1.57 dB for *w*_0_ and −0.99 to −1.07 dB for *p* (Sec. S2).

## 4. Discussion

### 4.1 Model adequacy, peripheral compression, and read-out comparison

Across 250 Hz to 4 kHz, results support pairing peripheral compression (Eq. (1)) with across-channel suppression. M0 and the five-way ablation align with Hots *et al*. (2016), whose 1500-Hz dual-resonance nonlinear (DRNL) simulations (5- to 135-Hz noise) yielded 0.5 to 1 dB of compression reduction (2 dB with two-tone suppression), pointing to retrocochlear mechanisms. Likewise, multi-channel M0 yields −0.17 dB for 1/4-octave noise (+5.44 dB for 1-octave noise) because off-frequency summation cancels within-channel compression, whereas 20-Hz SAM inside one filter yields −2.28 dB (+0.20 dB under RMS pre-compression); suppression unmasks compressive depression in noise.

Rather than identifying a biological circuit, the benchmark separates structures across two criteria. On ≤ 1-octave CLS data, all five Class 3 models perform indistinguishably near the 1.68-dB SE reference (48-curve and 20-listener bootstraps), with global NEA (Eq. (2)) ranking best by curve-level BIC_48_ (point-level *n*_eff_ = 380 favors six-parameter hybrids) and CLS transfer. Local divisive normalization (Eq. (3)) yields the lowest zero-refit error on the single external broadband dataset (*N* = 7; Leibold *et al*., 2007), beating global NEA while overlapping local sub-WBI under CLS parameter uncertainty. The two read-outs are closely related: for a kernel wider than the activity pattern, Eq. (3) is the summed activity divided by the (*p* − 1)th power of an equivalent width of the pattern, and Eq. (2) is the same with the width taken from the maximum and the power *w*; they differ mainly in the finite span of the kernel, which only stimuli wider than the span can test (Sec. S2). Its Gaussian SD (*σ*_ERB_ = 3.40 ERBs; CI [2.58, 5.39]) spans 2.16 octaves (±1*σ*_ERB_) at 250 Hz, 1.31 at 1 kHz, and 1.12 at 4 kHz. While local shunting pooling aligns with ventral cochlear nucleus D-stellate inhibition onto T-stellate principal cells (Carandini and Heeger, 2012; Oertel *et al*., 2011; Palmer *et al*., 1996; Pressnitzer *et al*., 2001; Winter and Palmer, 1995), aggregate behavioral fits cannot identify the neural circuit.

### 4.2 Substantive limitations and clinical outlook

Four limitations remain open. First, both models track mean levels well but miss fine structure on subsets (Eq. (2) misses low-level noise recovery; Eq. (3) yields weak/negative SAM correlations, Fig. 3) and underestimate the 1/4-octave noise dip; level-dependent filter broadening (Glasberg and Moore, 2000) and dynamic modulation models remain open. Second, flat noise and SAM tones came from separate cohorts differing in age, screening, and trial count (200 vs. 100 trials without practice), contributing to the 3.85-CU high-level offset and transfer asymmetry; external broadband testing rests on one six-point dataset (*N* = 7; Leibold *et al*., 2007), motivating within-listener noise, SAM, and broadband data. Third, because SuppPub2.xlsx pairs 30-listener BTNRH group maps with 20-listener UW boundaries (trial-by-trial BTNRH data are archived in Neely, 2026), joint CIs reflect 48-curve cluster uncertainty alongside exact 20-listener UW resampling. Fourth, conclusions apply strictly to 250 Hz to 4 kHz (excluding the two 1-octave curves at 4 kHz or applying within-band ER-3A filtering across 250 Hz to 4 kHz left RMSEs within 0.02 dB and all rankings unchanged; Table S2); higher frequencies require inverse-filter equalization (Sec. S3).

Both *w*_0_ (one central parameter, no fixed pooling constants) and *p* (decoupled from threshold tone growth via *s*_0_ and restoring summation above 1 octave) offer compact candidates for individualized loudness profiling, shifting 1-octave matches by 1.0 to 1.6 dB per 0.1 unit. Hots *et al*. (2016) observed roughly 3.5 dB mean mid-bandwidth depression (with wide variability) in hearing-impaired listeners despite reduced cochlear compression, motivating identifiability tests of both read-outs in hearing-impaired ears (250 Hz to 4 kHz).

## 5. Summary

Across 250 Hz to 4 kHz, peripheral compression alone fails on wideband noise, whereas pairing a peripheral stage with a memoryless compression nonlinearity and across-channel suppression captures normal-hearing CLS bandwidth, level, and AM effects near the 1.68-dB listener SE reference. Curve-level CLS parsimony (*n*_eff_ = 48) favors global NEA, whereas local divisive normalization yields the lowest observed zero-refit error on the single external broadband dataset (*N* = 7).

## Supplementary Material

See SuppPub1 for peripheral and nine-model parameters (Tables S1a and S1b), benchmark, 4-kHz/ER-3A sensitivity, and five-level BIC_eff_ tables (Tables S2 and S3), ablations (Table S4), *B* = 1, 000 UW and Leibold CIs (Table S5), equivalent-width derivation and exploratory 6-kHz analyses (Secs. S2 and S3, Fig. S1), dataset (`SuppPub2.xlsx`), and standalone scripts (`SuppPub3.py, SuppPub4.py`).

## Acknowledgments

Supported by NIH Grant Nos. R01 DC017988 (PI: Shen) and R01 DC008318 (PI: Neely), and the Mary Gates Undergraduate Research Scholarship (Shohet). The authors thank Erik A. Petersen, Calista Shohet, Brianna Chessik, Nicolas McArthur, Sara E. Harris, Joshua J. Hajicek, and Daniel Rasetshwane for data collection and qCLS development. Large language models (Gemini, Google; Claude, Anthropic) assisted with coding and drafting; the authors verified the code, the analyses, and the text.

## Author Declarations

*Conflict of Interest.* The authors have no conflicts to disclose. *Ethics Approval.* Experimental protocols were approved by the Institutional Review Boards of the University of Washington and Boys Town National Research Hospital, and all participants provided written informed consent. *Author ORCIDs.* Yi Shen: https://orcid.org/0000-0003-0063-1200; Stephen T. Neely: https://orcid.org/0000-0002-1349-5386.

## Data Availability

BTNRH trial-by-trial CLS data are available at https://doi.org/10.5281/zenodo.23090549 (Neely, 2026). The integrated dataset including de-identified UW category boundaries (`SuppPub2.xlsx`) and standalone Python scripts for primary analyses (`SuppPub3.py`) and diagnostics (`SuppPub4.py`) are published as supplementary material.

## Notes

### Competing Interest Statement

The authors have declared no competing interest.

## References and links

Carandini, M., and Heeger, D. J. (2012). “Normalization as a canonical neural computation,” Nat. Rev. Neurosci. 13, 51–62.

Dau, T., Kollmeier, B., and Kohlrausch, A. (1997). “Modeling auditory processing of amplitude modulation. I. Detection and masking with narrow-band carriers,” J. Acoust. Soc. Am. 102, 2892–2905.

Glasberg, B. R., and Moore, B. C. J. (1990). “Derivation of auditory filter shapes from notched-noise data,” Hear. Res. 47, 103–138.

Glasberg, B. R., and Moore, B. C. J. (2000). “Frequency selectivity as a function of level and frequency measured with uniformly exciting notched noise,” J. Acoust. Soc. Am. 108, 2318–2328.

Glasberg, B. R., and Moore, B. C. J. (2002). “A model of loudness applicable to time-varying sounds,” J. Audio Eng. Soc. 50, 331–342.

Hots, J., Jarzombek, K., and Verhey, J. L. (2016). “Mid-bandwidth loudness depression in hearing-impaired listeners,” J. Acoust. Soc. Am. 139, 2334–2341.

Hots, J., Rennies, J., and Verhey, J. L. (2014). “Loudness of subcritical sounds as a function of bandwidth, center frequency, and level,” J. Acoust. Soc. Am. 135, 1313–1320.

Jepsen, M. L., Ewert, S. D., and Dau, T. (2008). “A computational model of human auditory signal processing and perception,” J. Acoust. Soc. Am. 124, 422–438.

Leibold, L. J., Tan, H., Khaddam, S., and Jesteadt, W. (2007). “Contributions of individual components to the overall loudness of a multitone complex,” J. Acoust. Soc. Am. 121, 2822–2831.

Moore, B. C. J., Glasberg, B. R., and Baer, T. (1997). “A model for the prediction of thresholds, loudness, and partial loudness,” J. Audio Eng. Soc. 45, 224–240.

Neely, S. T. (2026). “Quick tests of consonant confusion and loudness growth,” Zenodo doi: 10.5281/zenodo.23090549.

Neely, S. T., Harris, S. E., Hajicek, J. J., Petersen, E. A., and Shen, Y. (2026). “Modeling the influence of bandwidth and envelope on categorical loudness scaling,” bioRxiv 2026.03.30.715393, doi: 10.64898/2026.03.30.715393.

Neely, S. T., and Jesteadt, W. (2005). “Quadratic-compression model of auditory discrimination and detection,” Acta Acust. united Ac. 91, 980–991.

Oertel, D., Wright, S., Cao, X.-J., Ferragamo, M., and Bal, R. (2011). “The multiple functions of T stellate/multipolar/chopper cells in the ventral cochlear nucleus,” Hear. Res. 276, 61–69.

Palmer, A. R., Jiang, D., and Marshall, D. H. (1996). “Responses of ventral cochlear nucleus onset and chopper units as a function of signal bandwidth,” J. Neurophysiol. 75, 780–794.

Pressnitzer, D., Meddis, R., Delahaye, R., and Winter, I. M. (2001). “Physiological correlates of comodulation masking release in the mammalian ventral cochlear nucleus,” J. Neurosci. 21, 6377–6386.

Rabinowitz, N. C., Willmore, B. D. B., Schnupp, J. W. H., and King, A. J. (2011). “Contrast gain control in auditory cortex,” Neuron 70, 1178–1191.

Shen, Y., Petersen, E. A., and Neely, S. T. (2024). “Toward parametric Bayesian adaptive procedures for multi-frequency categorical loudness scaling,” J. Acoust. Soc. Am. 156, 262–277.

Winter, I. M., and Palmer, A. R. (1995). “Level dependence of cochlear nucleus onset unit responses and facilitation by second tones or broadband noise,” J. Neurophysiol. 73, 141–159.

Zwicker, E., and Scharf, B. (1965). “A model of loudness summation,” Psychol. Rev. 72, 3–26.

