## Supplemental tables for "Compressive peripheral processing and across-channel suppression models of loudness bandwidth effects"

Provides complete peripheral and nine-model central parameters using 1-dB log-loudness tone-growth inversion (Tables S1a and S1b), the nine-model benchmark with 4-kHz / within-band ER-3A sensitivity rows and five-level BIC sensitivity (Tables S2 and S3), five-way peripheral ablations (Table S4), B = 1,000 exact 20-listener UW bootstrap and 48-curve Leibold parameter-uncertainty CIs (Table S5), equivalent-width derivation between Eqs. (2) and (3), and exploratory 6-kHz calibration analyses (Sec. S3 and Fig. S1). Companion files: SuppPub2.xlsx (clean 7-sheet workbook), SuppPub3.py, and SuppPub4.py.

**S1. Complete peripheral and nine-model central parameter specifications**

Table S1a. Shared 13-parameter pure-tone peripheral stage with a memoryless compression nonlinearity (Eq. 1; 250 Hz to 4 kHz).

| **Parameter (units)** | **Fitted value** | **Constraint** |
| --- | --- | --- |
| Gain cubic g0, g1, g2, g3 | -0.0194, +0.1488, -0.2536, +0.0499 | BTNRH pure-tone growth (250 to 4000 Hz) |
| Compression a0, a1 | 0.8761, +0.0654 | BTNRH pure-tone growth |
| Compression b0, b1 | 0.01181, -0.00260 | BTNRH pure-tone growth |
| Compression c0, c1 | -1.22e-07, +1.36e-07 | BTNRH pure-tone growth |
| Rate exponent alpha / knee I0 | 0.4681 / 18.48 (12.7 dB) | BTNRH pure-tone growth |
| Loudness constant C | 0.0922 | Pure-tone growth (13.4%, 1.85 dB) |

Table S1b. Fitted central parameters and optimization bounds for all nine models (250 Hz to 4 kHz).

| **Model** | **k** | **Fitted parameters (250 Hz to 4 kHz)** | **Optimization bounds** |
| --- | --- | --- | --- |
| M0: Classical sum | 0 | None | None |
| M3a: 12-band MFB only | 3 | kappa=0.74, f_{m,c=400.0 Hz, beta=-0.13 | [0,8], [20,400], [-0.8,0.8] |
| M2a: Local sub-WBI (linear) | 2 | gamma=0.154, sigma_{mathrm{ERB=2.61 ERBs | [0,0.35], [1.5,50] |
| M2b: Local sub-WBI (sat.) | 4 | gamma=0.161, sigma=2.53, log_{10E_{mathrm{th=+0.20, log_{10E_{mathrm{sat=4.90 | [0,0.5], [1.5,50], [-2,3], [0.5,5] |
| M1a: Global NEA (const.) | 1 | w_0=0.636 [0.61, 0.68] | [0,1.3] |
| M1b: Gated NEA (activity) | 3 | w_0=0.666, mu_0=2.85, sigma_w=3.35 | [0,1.3], [-0.5,4], [0.3,5] |
| M2c: Divisive norm (Eq. 3) | 3 | p=2.129 [1.93, 2.37], sigma_{mathrm{ERB=3.40 [2.58, 5.39], s_0=4.06 [2.76, 5.33] | [1,3], [1.5,40], log_{10}s_0in[-2,3] |
| M4b: 12-band MFB + Eq. 3 | 6 | kappa=0.29, f_{m,c=397, beta=-0.46, p=1.93, sigma=5.97, log_{10s_0=+0.96 | Union of M3a and M2c bounds |
| M5: 12-band MFB + NEA | 6 | kappa=0.19, f_{m,c=393, beta=-0.62, w_0=0.66, mu_0=2.95, sigma_w=2.20 | Union of M3a and M1b bounds |

**S2. Benchmark, 4-kHz / ER-3A sensitivity, BIC sensitivity, ablations, and bootstraps**

Table S2. Nine-model benchmark across 380 valid CLS observations (250 Hz to 4 kHz) plus 4-kHz 1-oct exclusion and within-band ER-3A sensitivity rows.

| **Model** | **k** | **r** | **Shared (dB)** | **Match (dB)** | **BIC_48** | **BT to UW (dB)** | **UW to BT (dB)** | **Leib0 (dB)** | **Leib1 (dB)** |
| --- | --- | --- | --- | --- | --- | --- | --- | --- | --- |
| M0: Classical sum | 0 | +0.504 | 4.34 | 5.11 | 140.9 | 3.33 | 5.90 | 4.62 | 4.62 |
| M3a: 12-band MFB only | 3 | +0.578 | 3.59 | 4.50 | 134.3 | 5.02 | 5.31 | 3.70 | 3.70 |
| M2a: Local sub-WBI (linear) | 2 | +0.598 | 1.76 | 1.76 | 62.2 | 2.23 | 2.26 | 2.42 | 2.02 |
| M2b: Local sub-WBI (sat.) | 4 | +0.628 | 1.73 | 1.73 | 68.2 | 2.28 | 2.26 | 2.46 | 2.03 |
| M1a: Global NEA (const.) | 1 | +0.647 | 1.63 | 1.63 | 51.0 | 1.53 | 1.93 | 3.37 | 1.79 |
| M1b: Gated NEA (activity) | 3 | +0.664 | 1.61 | 1.60 | 57.1 | 1.70 | 1.93 | 3.73 | 1.81 |
| M2c: Divisive norm (Eq. 3) | 3 | +0.623 | 1.67 | 1.67 | 60.8 | 1.98 | 2.35 | 1.90 | 1.48 |
| M4b: 12-band MFB + Eq. 3 | 6 | +0.704 | 1.50 | 1.50 | 62.3 | 2.00 | 2.41 | 2.29 | 1.33 |
| M5: 12-band MFB + NEA | 6 | +0.723 | 1.47 | 1.49 | 60.5 | 2.04 | 2.21 | 3.61 | 1.56 |
| M1a (excl. 4-kHz 1-oct, n=365) | 1 | +0.630 | 1.63 | n/a | n/a | n/a | n/a | 3.47 | n/a |
| M2c (excl. 4-kHz 1-oct, n=365) | 3 | +0.608 | 1.66 | n/a | n/a | n/a | n/a | 2.09 | n/a |
| M1a (within-band ER-3A, n=380) | 1 | +0.640 | 1.65 | n/a | n/a | n/a | n/a | 3.33 | n/a |
| M2c (within-band ER-3A, n=380) | 3 | +0.633 | 1.65 | n/a | n/a | n/a | n/a | 1.84 | n/a |

Table S3. Matched peripheries, five-level BIC sensitivity (n_eff = 12, 24, 30, 48, 380), and 48-curve cluster-bootstrap comparisons against M2c.

| **Model** | **k** | **Shared** | **Match** | **BIC_12** | **BIC_24** | **BIC_30** | **BIC_48** | **BIC_380** | **Diff vs. M2c [95% CI]** | **Prop(M2c)** |
| --- | --- | --- | --- | --- | --- | --- | --- | --- | --- | --- |
| M0: Classical sum | 0 | 4.34 | 5.11 | 35.2 | 70.5 | 88.1 | 140.9 | 1115.7 | +2.67 [+1.91, +3.40] | 1.000 |
| M3a: 12-band MFB only | 3 | 3.59 | 4.50 | 38.1 | 70.9 | 86.9 | 134.3 | 989.4 | +1.92 [+1.34, +2.49] | 1.000 |
| M2a: Local sub-WBI (linear) | 2 | 1.76 | 1.76 | 18.6 | 33.6 | 40.8 | 62.2 | 442.8 | +0.09 [-0.04, +0.21] | 0.926 |
| M2b: Local sub-WBI (sat.) | 4 | 1.73 | 1.73 | 23.1 | 39.1 | 46.6 | 68.2 | 441.4 | +0.06 [-0.06, +0.18] | 0.851 |
| M1a: Global NEA (const.) | 1 | 1.63 | 1.63 | 14.3 | 26.7 | 32.8 | 51.0 | 378.7 | -0.04 [-0.14, +0.06] | 0.240 |
| M1b: Gated NEA (activity) | 3 | 1.61 | 1.60 | 18.8 | 32.3 | 38.6 | 57.1 | 378.0 | -0.06 [-0.17, +0.03] | 0.097 |
| M2c: Divisive norm (Eq. 3) | 3 | 1.67 | 1.67 | 19.8 | 34.1 | 41.0 | 60.8 | 407.5 | Ref. | Ref. |
| M4b: 12-band MFB + Eq. 3 | 6 | 1.50 | 1.50 | 24.7 | 38.6 | 44.9 | 62.3 | 345.4 | -0.17 [-0.27, -0.06] | 0.001 |
| M5: 12-band MFB + NEA | 6 | 1.47 | 1.49 | 24.2 | 37.7 | 43.7 | 60.5 | 330.8 | -0.20 [-0.32, -0.05] | 0.004 |

Table S4. Five-way peripheral compression ablation under M0 and M2c (primary global-DE M2c fit for Full Eq. 1: 1.90 dB Leibold RMSE).

| **Peripheral variant** | **M0 RMSE** | **M0 20Hz** | **M0 100Hz>=1.5k** | **M0 BTq** | **M0 BT1** | **M2c RMSE** | **M2c 20Hz** | **M2c Leib0** |
| --- | --- | --- | --- | --- | --- | --- | --- | --- |
| Full Eq. (1) (primary fit) | 4.33 | -2.28 | -1.41 | -0.17 | +5.44 | 1.67 | -2.64 | 1.90 |
| RMS envelope before Eq. (1) | 5.31 | +0.20 | +0.56 | +1.30 | +7.01 | 2.43 | -0.01 | 3.00 |
| Rate nonlinearity only (E = L_{mathrm{env) | 3.16 | -1.43 | -0.94 | -0.18 | +3.19 | 1.80 | -1.64 | 2.12 |
| Quadratic E_k only (linear rate) | 2.77 | -0.88 | -0.60 | -0.17 | +1.85 | 1.99 | -1.07 | 3.14 |
| Linear energy (bar{s_k propto e_{mathrm{rms^2) | 2.63 | +0.09 | -0.00 | -0.01 | +0.06 | 2.44 | +0.02 | 4.62 |

Table S5. Exact 20-listener UW bootstrap (B = 1,000) and 48-curve Leibold CLS parameter-uncertainty propagation (B = 1,000).

| **Model** | **UW 20-list. RMSE [95% CI]** | **UW diff vs M2c [95% CI]** | **Leib0 obs.** | **Leib0 CLS-param CI** | **Leib diff vs M2c [95% CI]** | **Prop(M2c)** |
| --- | --- | --- | --- | --- | --- | --- |
| M0: Classical sum | 3.66 [2.75, 4.78] | +1.61 [+0.83, +2.38] | 4.62 | [4.62, 4.62] | [+1.81, +2.95] | 1.000 |
| M2a: Local sub-WBI | 2.09 [1.42, 3.21] | +0.08 [-0.15, +0.36] | 2.42 | [2.23, 2.84] | [-0.31, +0.76] | 0.929 |
| M2b: Sat. sub-WBI | 2.07 [1.40, 3.19] | +0.05 [-0.15, +0.33] | 2.46 | [2.29, 2.92] | [-0.30, +0.83] | 0.931 |
| M1a: Global NEA | 2.08 [1.45, 3.18] | +0.07 [-0.11, +0.30] | 3.37 | [3.06, 3.77] | [+0.45, +1.89] | 0.991 |
| M1b: Gated NEA | 2.05 [1.43, 3.09] | +0.02 [-0.11, +0.17] | 3.73 | [3.32, 4.18] | [+0.83, +2.22] | 0.996 |
| M2c: Divisive norm | 2.03 [1.43, 3.06] | Ref. | 1.90 | [1.67, 2.81] | Ref. | Ref. |

**S3. Exploratory 6-kHz analysis**

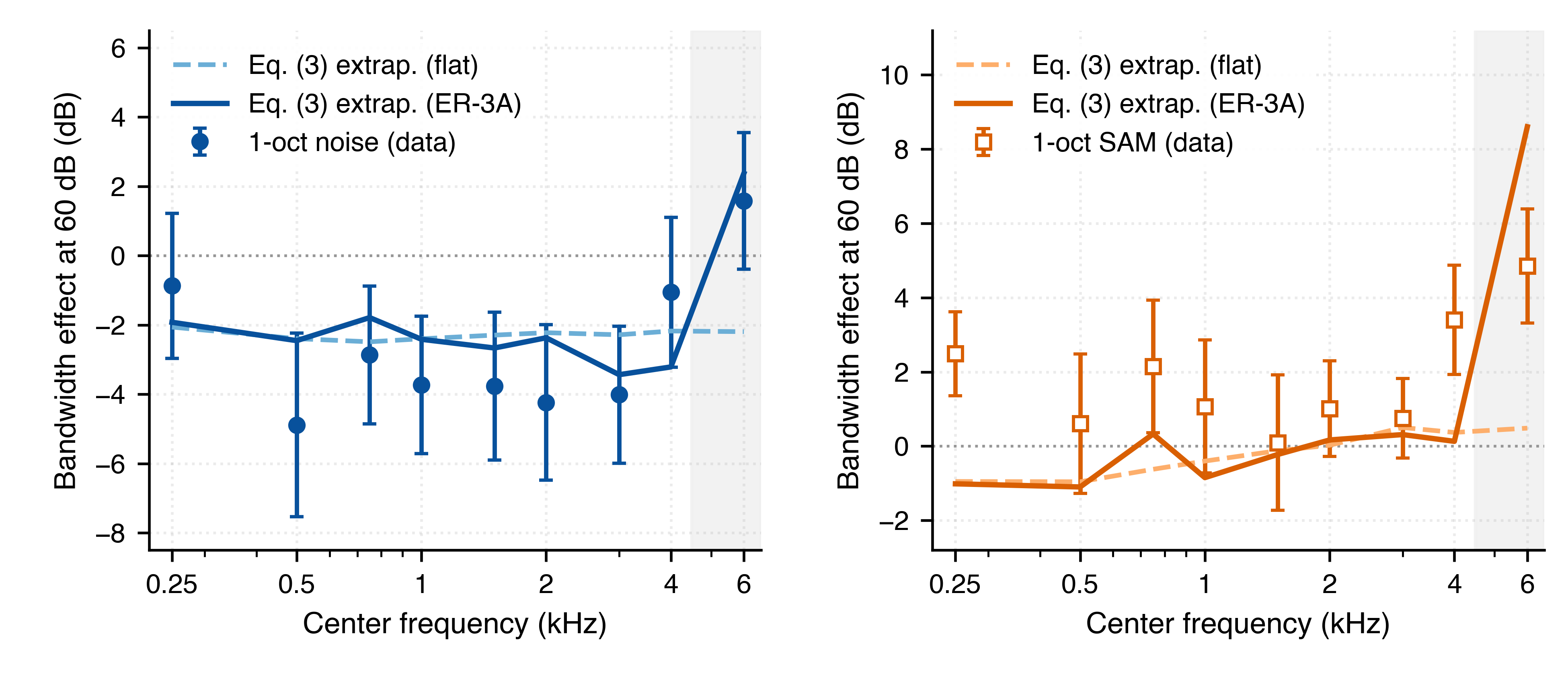

Figure S1. Exploratory zero-refit extrapolation of Eq. (3) to 6 kHz at 60 dB SPL.
